# Vascular tortuosity quantification as an outcome metric of the oxygen-induced retinopathy model of ischemic retinopathy

**DOI:** 10.1101/2022.10.02.510568

**Authors:** Kyle V. Marra, Jimmy S. Chen, Hailey K. Robles-Holmes, Joseph Miller, Guoqin Wei, Edith Aguilar, Yoichiro Ideguchi, Kristine B. Ly, Sofia Prenner, Deniz Erdogmus, Napoleone Ferrara, J. Peter Campbell, Martin Friedlander, Eric Nudleman

**Author notes:** Corresponding Author: Eric Nudleman, MD, PhD, Viterbi Family Department of Ophthalmology and Shiley Eye Institute, University of California San Diego, 9415 Campus Point Dr, MC 0946, La Jolla, CA 92093-0946, USA. Kyle V. Marra and Jimmy S Chen contributed to this work equally.

## Abstract

The murine oxygen-induced retinopathy (OIR) model is one of the most widely used animal models of ischemic retinopathy, mimicking hallmark pathophysiology of initial vaso-obliteration (VO) resulting in ischemia that drives neovascularization (NV). In addition to NV and VO, human ischemic retinopathies including Retinopathy of Prematurity (ROP) are characterized by increased vascular tortuosity. Vascular tortuosity is an indicator of disease severity, need to treat, and treatment response in ROP. Current literature investigating novel therapeutics in the OIR model report their effects on NV and VO, but no standardized quantification of vascular tortuosity exists to date despite this metric’s relevance to human disease in clinics. The current proof-of-concept study applied a computer-based image analysis algorithm capable of calculating standardized measurements of vascular tortuosity. Quantification of vascular tortuosity correlated with disease activity in OIR analogously to that observed in infants with ROP. Treatment of OIR mice with anti-Vascular Endothelial Growth Factor (aflibercept) rescued vascular tortuosity in the model. Altogether, these data demonstrated that vascular tortuosity is a quantifiable feature of the OIR model and may be used as an outcome measurement in future studies investigating new treatment modalities for retinal ischemia.

## INTRODUCTION

Ischemic retinopathies such as diabetic retinopathy (DR), retinopathy of prematurity (ROP), and retinal vein occlusions remain leading causes of blindness in the world.(1, 2) With over 1,000 publications on the model since its inception in 1994, the murine oxygen-induced retinopathy (OIR) model is one of the most widely-utilized animal models in preclinical studies aimed at developing novel therapeutic strategies for ischemic retinopathies.(3) OIR mice recapitulate key features of ischemic retinopathy – initial vaso-obliteration (VO) followed by neovascularization (NV) – and additional sequelae such as vascular leakage and proliferation.(4, 5) Ischemic retinopathies like ROP are characterized by aberrant neovascularization and avascular zones, however, these conditions also present with increased vascular dilation and tortuosity. Tortuosity is a crucial indicator of disease severity, need to treat, and treatment response in ROP.(6) Preclinical research in OIR mice typically report the effect of therapeutic interventions on rescuing NV and VO, and rarely comment on changes to vascular tortuosity. In fact, a standardized methodology to calculate vascular tortuosity in OIR mice does not currently exist and the effects of OIR on vascular tortuosity remain uncharacterized.

The current proof-of-concept study applied a previously published semi-automated computer-based image analysis approach towards investigating the correlation between retinal vascular tortuosity and disease activity in OIR mice analogous to that observed in infants with ROP. This study also investigated whether treatment of OIR mice rescues vascular tortuosity. Vascular studies using deep convolutional neural networks to quantify tortuosity in human retinal images of ROP can now outperform human experts.(7) Using the outputs of this deep learning model, recent studies have been able to develop a quantitative metric of vascular tortuosity using an algorithm called iROP-Assist that takes into account features like vascular dilation and tortuosity.(8) Such a metric does not exist to quantify tortuosity in OIR mice, the classic model for ROP. In the current study, we utilized the capabilities of iROP-Assist to generate a cumulative tortuosity index (CTI) measurement that may be used as a novel outcome measurement in future experiments using the OIR model. Manually segmented OIR images were analyzed with the iROP-Assist algorithm to establish which measurements of tortuosity had the best discrimination between OIR mice and wild-type controls. The degree of retinal vascular tortuosity in OIR mice correlated with the various stages of the model’s pathophysiology. Agents known to rescue NV in the OIR phenotype (aflibercept) were found to rescue CTI in OIR mice. Altogether, this work provides a new tool to quantify vascular tortuosity in OIR mice and establishes CTI as a meaningful outcome measurement in future preclinical studies aimed at developing the next armamentarium of treatment agents for ROP and other ischemic retinopathies.

## RESULTS

### Cross-validation of manual segmentations

To ensure that manual segmentation of tortuosity was reliably quantified between graders, a cross-validation experiment was performed on a sample of randomly selected age-matched retinas of mice sacrificed at P12, P17, and P25 under normoxic (NOX) and OIR conditions from a previously published dataset.(5) Manual vessel segmentation was performed by four graders (JSC, HKB, SP, and JM). Cross-validation of manual segmentations was achieved by comparing 20 flat mounted retinas (10 NOX and 10 OIR) from the age-matched experimental dataset. All vessel segmentations were reviewed by two independent graders (JSC and JM) to verify that selection of large vessels for segmentation was similar among all graders. For each image, all vessel segmentations produced by each grader were compared quantitatively using the Dice coefficient. Comparison of the Dice coefficient for all 6 combinations of 4 graders (0.79 ± 0. 13, 0.58 ± 0.27, 0.63 ± 0. 25, 0.61 ± 0.1, 0.61 ± 0.21, and 0.57 ± 0.24) demonstrated no significant differences in vascular segmentation (p=0.24), suggesting manual segmentation was a reliable method for future experiments. Representative images of flat-mounted retinas and corresponding segmentations among all 4 graders are shown in **Figure 1**.

**Figure 1.**
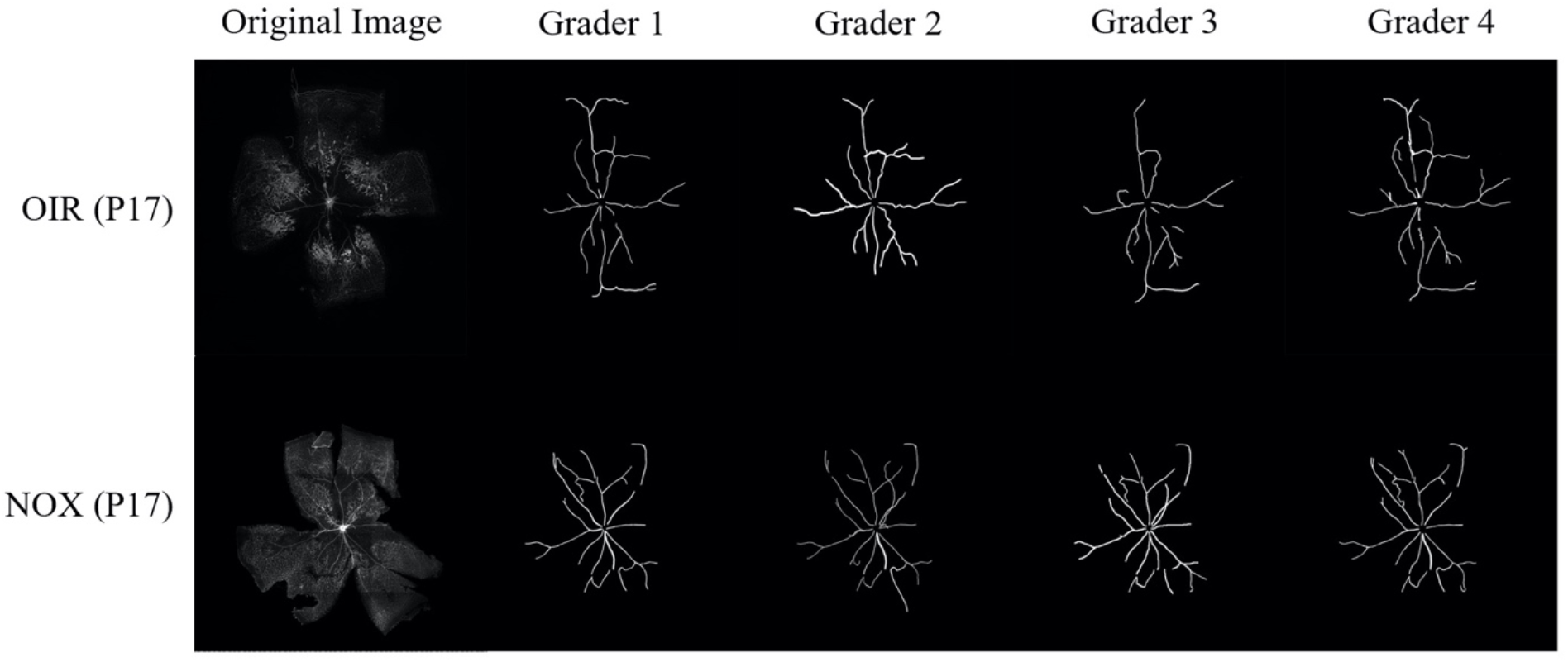
Representative segmentations of flat-mounted retinas of NOX and OIR mice from 4 independent graders. OIR (top) and NOX (bottom) mice sacrificed at P17 were stained with GS-IB4 and flat-mounted for manual quantification by 4 independent graders. All segmentations were reviewed by an expert ophthalmologist. Following expert review of image segmentation from an ophthalmologist (JC), Dice coefficients were calculated to compare the similarity of vessel segmentations between all graders.

### Pilot assessment of vascular tortuosity metrics

A pilot study was performed to investigate metrics of tortuosity including the integrated curvature (IC), overall curvature (OC), and cumulative tortuosity index (CTI) on NOX versus OIR retinas from the same dataset.(5) These metrics were derived from a previously published artificial intelligence algorithm that calculates various metrics to assess plus disease in ROP.(8) As described by Boton-Canedo et al.(9) and Han (10), CTI measures the mean cumulative sums of angles between segmented vessels normalized by vessel length, OC measures the mean angle curvature for all segments, and the IC is the integral of the OC. All vascular tortuosity metrics demonstrated statistically significant differences (p < 0.0001) when discriminating vascular tortuosity between NOX and OIR mice (**Figure 2**). CTI was selected for use in all subsequent experiments since its methodology is the most intuitive for measuring vascular tortuosity.

**Figure 2.**
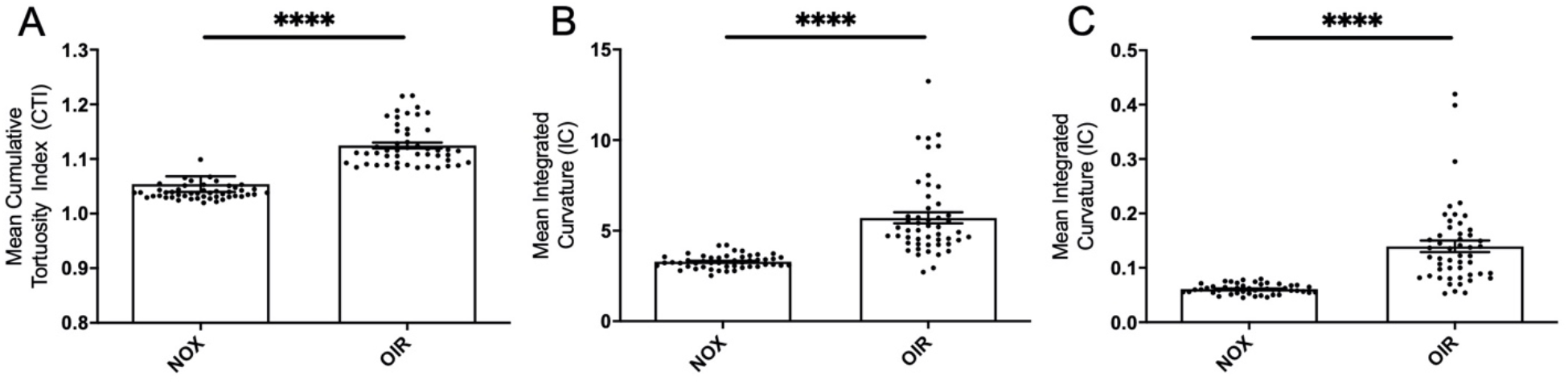
Quantification of vascular tortuosity in NOX and OIR mice using previously defined metrics. Manual segmentation of 50 NOX images and 50 OIR retinal images from mice sacrificed at P17 were quantified using a published algorithm that extracts various tortuosity metrics including (**A**) cumulative tortuosity index, (**B**) integrated curvature, and (**C**) overall curvature. Two-tailed Student’s *t* test demonstrated statistically significant differences between all vascular tortuosity metrics when comparing images from NOX and OIR mice. Results were compared using a two-tailed Student’s *t* test; error bars represent SEM; ****p < 0.0001

### Assessment of vascular tortuosity between age-matched NOX vs OIR images

To investigate the validity of quantifying vascular tortuosity as an outcome measurement of OIR mice, manually segmented flat mount images from age-matched NOX and OIR mice were analyzed at timepoints of pathophysiological relevance in the model. Mice placed in hyperbaric oxygen on P7 were sacrificed on P12 immediately upon return to room air at the onset of the ischemic drive, on P17 at the peak of the neovascular phase, and on P25 when the retinal vasculature architecture is partially restored. These retinal images were obtained from a previous dataset(5) with additional numbers obtained using the same methodology and by the same authors. The total number of images analyzed at each timepoint is shown in **Table 1**.

**Table 1.** Number of normoxic and OIR mice used in experiments characterizing neovascularization, vaso-obliteration, and cumulative tortuosity index.

| **Age** | **<u>Number of Retinas (n)</u>** |  |
| --- | --- | --- |
|  | **Normoxic mice** | **OIR mice** |
| P12 | 42 | 34 |
| P17 | 37 | 50 |
| P25 | 48 | 32 |

The mean ± SD NV ratio for NOX images at all timepoints was 0, and the mean ± SD NV ratio for OIR images was 0.01 ± 0.01 at P12 (p = 0.01), 0.08 ± 0.04 at P17 (p < 0.0001), and 0.03 ± 0.02 at P25 (p < 0.0001) (**Figure 3A)**. The mean ± SD VO ratio for NOX and OIR images was 0.09 ± 0.04 vs. 0.26 ± 0.06 at P12 (p < 0.0001), 0.11 ± 0.04 vs. 0.15 ± 0.07 at P17 (p = 0.004), and 0.08 ± 0.06 vs 0.05 ± 0.02 at P25 (p = 0.004) (**Figure 3B**). The mean ± SD CTI for NOX and OIR images was 1.04 ± 0.02 vs. 1.04 ± 0.02 (p = 0.6) at P12, 1.05 ± 0.01 vs. 1.14 ± 0.09 (p < 0.0001) at P17, and 1.04 ± 0.01 vs. 1.08 ± 0.04 respectively (p < 0.0001) at P25 **(Figure 3C)**. Together, these data suggest that vascular tortuosity is a valid and distinguishable feature of the OIR model at P17 and P25, time points of pathologic significance.

**Figure 3.**
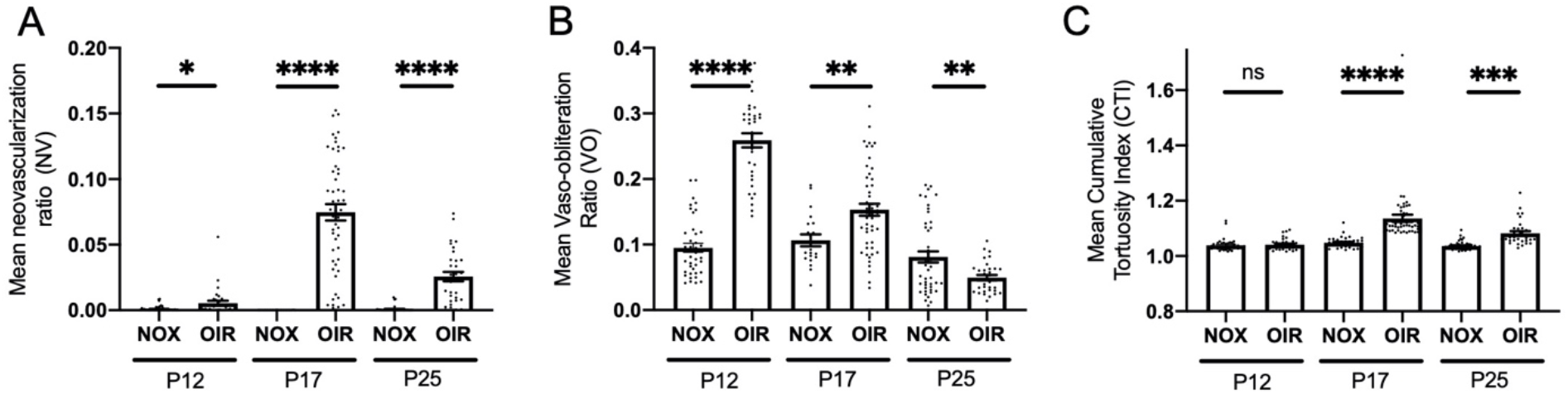
Characterization of neovascularization, vaso-obliteration, and vascular tortuosity in age-matched NOX and OIR retinas at P12, P17, and P25. (**A-C**) NV, VO, and vascular tortuosity measured as CTI demonstrated changes congruent with the pathophysiology of OIR. (**A-B**) Quantified using a publicly available deep learning algorithm, the (**A**) NV and (**B**) VO of NOX and OIR mice were consistent with previous reports.(5) In OIR mice, NV peaks at P17 and is partially resolved by P25 while VO peaks at P12 and subsequently decreased over time. (**C**) Quantification of CTI using a previously published algorithm demonstrated that similar to NV, vascular tortuosity is increased at P17 and partially resolves by P25.(8) Results were compared using a two-tailed Student’s *t* test; error bars represent SEM; ns = non-significant, *p < 0.05, **p < 0.01, ***p < 0.001, ****p < 0.0001.

### Vascular tortuosity in mouse eyes treated with aflibercept vs IgG control

The next series of experiments were performed to investigate the effects of known therapeutic agents on vascular tortuosity in OIR mice using another previously published dataset.(11) Previous reports have demonstrated that intravitreal injections of aflibercept decrease neovascularization and retinal inflammation in OIR mice, but the effects of aflibercept on vascular tortuosity remain unknown.

Eyes of OIR mice placed in hyperbaric oxygen on P7 were intravitreally injected with aflibercept with contralateral eyes injected with IgG as control on P12 (immediately upon return to room air) and sacrificed on P17 for preparation of retinal flat mounts. A total of 36 images, 18 eyes treated with aflibercept and 18 treated with IgG control, were manually segmented and CTI was calculated using iROP-Assist. NV and VO ratios were calculated using previously published methods.(12) The mean ± SD NV ratio for aflibercept versus control was 0.01 ± 0.01 versus 0.03 ± 0.02 (p = 0.003) (**Figure 4A**). The mean ± SD VO ratio for aflibercept versus control was 0.10 ± 0.14 vs. 0.12 ± 0.05 (p = 0.6) (**Figure 4B**). Eyes treated with aflibercept demonstrated a mean ± SD CTI of 1.05 ± 0.04, whereas the control eyes demonstrated a mean CTI of 0.12 ± 0.06 (p = 0.002) (**Figure 4C**). Overall, these data suggest that anti-VEGF agents rescue the increased vascular tortuosity of the OIR model similar to the effect of anti-VEGF on vascular tortuosity in ROP.(13) Reduction in vascular tortuosity may serve as a useful quantitative outcome parameter in future experiments testing therapeutics in OIR mice.

**Figure 4.**
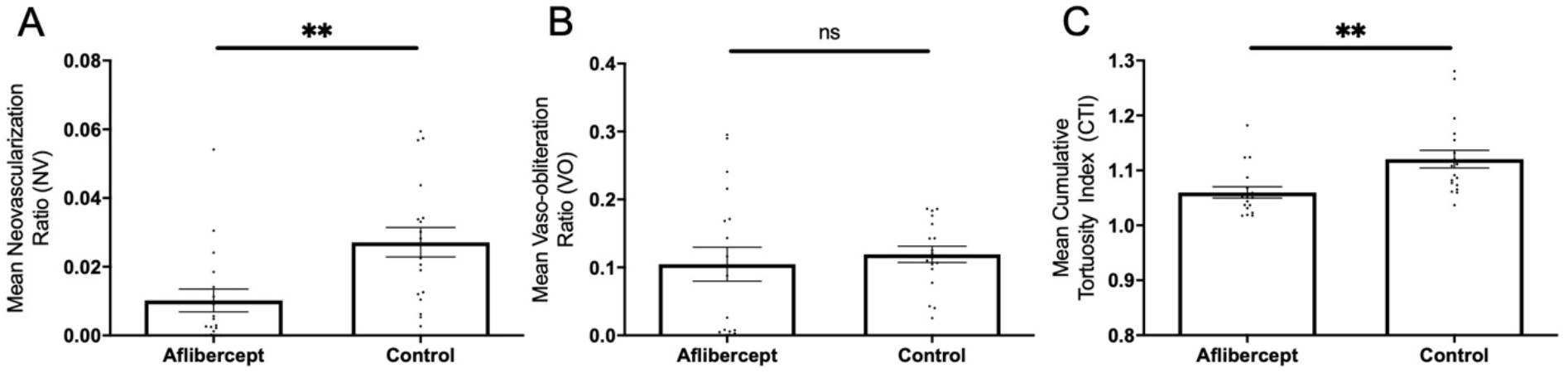
Characterization of neovascularization, vaso-obliteration, and vascular tortuosity in OIR mice following treatment with aflibercept. (**A-C**) Intravitreal injections of aflibercept improved retinal NV and CTI while the effect on VO did not reach statistical significance. A total of 18 images of eyes treated with aflibercept and 18 images of eyes treated with IgG control were manually segmented. (**A**) NV, (**B**) VO, and (**C**) CTI were calculated for all images and compared using a two-tailed Student’s *t* test; error bars represent SEM; ns = non-significant, *p < 0.05, **p < 0.01, ***p < 0.001, ****p < 0.0001.

## DISCUSSION

For years, the lack of a standardized definition of vascular tortuosity conditioned clinical specialists with subjective and thus varied conceptualizations of what constitutes tortuosity, leading to high inter- and even intra-expert variability in its quantification.(14, 15) Such ambiguity limited use of this metric as both a research and clinical tool. With the advent of explainable deep learning methodologies such as iROP-Assist, computational approaches capable of quantifying vascular tortuosity in humans have become clinically relevant.(6) In addition to its emerging role in the management of ROP, retinal vascular tortuosity is a clinical biomarker for numerous other vascular and systemic diseases including diabetic retinopathy, cerebrovascular disease, stroke, kidney dysfunction, and ischemic heart disease.(14, 16-19) The importance of vascular tortuosity to clinical medicine is an exciting and ongoing development. At the same time, the adaptation of these tools for use in preclinical studies on animal models of disease allows translational research to draw upon lessons learned from the clinics.

The current work, therefore, aimed to bring findings from the bedside back to the bench by accomplishing the following: 1.) adapting the iROP-Assist system to quantify vascular tortuosity in the retina of OIR and normoxic mice, 2.) characterizing the retinal vessel tortuosity at significant pathophysiologic time points in the OIR model, and 3.) demonstrating that therapeutic strategies used in the treatment of ROP have analogous effects in rescuing retinal vascular tortuosity in OIR mice. Altogether, these data suggest that vascular tortuosity in OIR mice is comparable to that seen in ROP and can be quantified using computer-based imaging analysis to provide a novel outcome measurement for experiments aimed at testing novel therapeutic agents *in vivo*.

The majority of literature using OIR mice report the retinal area covered by NV or VO on P17, typically in order to investigate phenotypic rescue following administration of a therapeutic agent on P12. Only a few studies have considered vascular tortuosity as an outcome measurement for the OIR model, and these publications report vascular tortuosity either by using an operationally defined scoring system or by calculating the ratio of tortuous to straight major vessels leaving the optic disc.(20-25). The current study is the first to provide a quantitative cumulative tortuosity index by using a computer-based image analysis algorithm that employs a feature-extraction-based approach to take into account all the major superficial vasculature within a retinal image. Analogously to NV, the CTI of OIR mice peaks at P17 and spontaneously regresses thereafter, as demonstrated by decreased CTI at P25. Intravitreal injection of the anti-VEGF agent aflibercept rescued both the NV as well as the CTI of OIR mice.

Altogether, this work applied techniques developed to quantify tortuosity in human ROP to its preclinical disease model of OIR. Analogously to ROP, OIR mice developed tortuosity that correlated with NV and successful treatment of the model rescued both outcome measurements. Given the clinical importance of assessing tortuosity in human ROP, measuring CTI in OIR mice may present a new and promising quantitative metric used in preclinical studies evaluating the efficacy of novel therapeutics in ROP.

## METHODS

### Animals

Age-matched C57BL/6 mice (The Jackson Laboratory, JAX) mice were subjected to normoxic conditions or to the OIR model as previously described.(3, 5) As described above, the first dataset was previously published(5) with additional mice prepared using the same methodology and by the same authors. In brief, pups were exposed to an atmosphere containing 75% oxygen from P7 to P12, after which they were returned to room air until euthanized at pre-specified time points: P12 (immediately), P17, and P25. Using previously published methods,(5) retinas were dissected from enucleated eyes by using fine brushes to separate and clean retina from choroid and sclera. Retinas were fixed in 4% paraformaldehyde on ice for 1 hour prior to overnight incubation in PBS with Ca^2+^Mg^2+^ with 10 µg *Griffonia simplicifolia-*Isolectin B4 (GS-IB4, I21412, Thermo Fisher Scientific). After being cut into 4 leaflets, retinas were flat-mounted for imaging using a Zeiss 710 confocal laser–scanning microscope with ZEN 2010 software (Zeiss) at 10x magnification and tile scanning (6 × 6 tiles). The second dataset was previously published by Xin et al. and consisted of pooled flat-mount retinal images from OIR mice intravitreally injected with aflibercept versus IgG contralaterally as control.(11)

### Manual segmentation and cross-validation

Flat-mount images of NOX and OIR retinas were manually segmented by 4 authors (JSC, HKRH, JM, and SP) for large vessels using Adobe Photoshop (Adobe; San Jose, CA). In summary, large vessels were defined as those that originated from the disc center as well as large vessel branches. Capillaries, neovascular tufts, and areas of vaso-obliteration were not segmented. To ensure segmentation methodology was similar between graders, cross-validation was performed on a set of 10 images from each group that were segmented by all authors. Combinations of all vessel segmentations for each image were first manually reviewed for similarity by two independent graders (JSC and JM). Vascular segmentation was then compared using Dice coefficient, which calculates the area of total overlap between the pixels segmented in 2 different images.

### Computer-based analysis of vascular tortuosity

A previously published algorithm for calculating plus disease on fundus images of infants with ROP was adapted to quantify vascular tortuosity of all segmented images and the associated coordinates of the optic disk center.(8) In summary, all unique pixels representing vessels from each segmentation were extracted to generate a graph of vessel segments. Point-based and vessel-based features such as integrated curvature, cumulative tortuosity index, and overall curvature were calculated from these vessel graphs using methods previously described by Ataer-Cansizoglu et al.(26) and definitions as described by Boton-Canedo et al.(9) and Han.(10) CTI measures the mean cumulative sums of angles between segmented vessels normalized by vessel length, OC measures the mean angle curvature for all segments, and the IC is the integral of the OC.

To investigate the validity of vascular tortuosity as a quantifiable feature of OIR, an initial pilot study using 20 randomly selected images (10 NOX and 10 OIR) from age-matched mice sacrificed at P12, P17, and P25 of a previously published dataset.(12) The mean IC, CTI, and OC were calculated for all 20 images and compared between groups. After analyzing the data, CTI was selected as the test metric of choice for all subsequent experiments quantifying vascular tortuosity since its methodology is the most appropriate for calculation of vascular tortuosity.

### Quantification of neovascularization and vaso-obliteration

Vaso-obliteration and neovascularization ratios were calculated as previously described.(12) In summary, a fully automated deep learning algorithm was used on images of retinal flat-mounts to generate segmentations for areas of NV and VO. The percentage of the images’ segmented area representing NV and VO were calculated as NV and VO ratios. These analyses were performed using a website hosting the deep learning system at http://oirseg.org/list.html.(12) All images for both experiments were input into this algorithm for generation of NV and VO ratios, and manually reviewed by a 2 graders (JSC and JM) to ensure that the segmentation accurately reflected pathology in the image. Of note, several NOX images were quantified by the deep learning algorithm to have high VO (likely due to areas of lighter pigmentation), but expert reviewers noticed that these images contained no vaso-obliteration as would be consistent with the natural physiology of vascular growth in normoxic mice. For these images, the VO was adjusted to agree with the experience of expert graders.

### Statistics

All statistical analyses were performed using R 4.0.5 (R Foundation, Vienna, Austria). For the pilot study and both experiments, mean CTIs, NV ratios, and VO ratios and their associated standard deviations were calculated at each time point for the first experiment (P12, P17, and P25) and were stratified by NOX vs OIR in both experiments. Statistical significance for experiments containing 2 groups was assessed using an unpaired Student’s *t* test with statistical significance threshold of p < 0.05. One-way ANOVA was used for comparison of 3 or more groups. Statistical significance was determined as *p < 0.05, **p < 0.01, ***p < 0.001, ****p < 0.0001 in all figures

### Study Approval

All experiments using animals were approved by Scripps Research Animal Care and Use Committee and the University of California at San Diego. Experiments were performed on C57BL/6 mice in accordance with the NIH Guide for the Care and Use of Laboratory Animals (National Academies Press, 2011). Protocols were approved by the Institutional Review Board at Scripps Research and Scripps Memorial Hospital, La Jolla, California, USA, and the University of California at San Diego. Study approval for previously published datasets was obtained as previously described.(5, 11)

## AUTHOR CONTRIBUTIONS

KVM and JC designed experiments, performed experiments, and wrote and edited the manuscript. HRH, JM, GW, EA, KL, and SP helped perform experiments. AI assisted in maintaining animal colonies in the lab of MF. NF and PC provided conceptual oversight. EN supervised the work and reviewed and edited the manuscript.

## ACKNOWLEDGEMENTS & FUNDING

We acknowledge and appreciate the many helpful discussions with members of The Scripps Research Institute and Lowy Medical Research Institute throughout the course of this project. This work was funded by grants to MF from the NIH National Eye Institute (NEI) (grant EY11254) and the Lowy Medical Research Institute. EN received funding from the NEI (K08 EY028999-01). KVM was funded by an F30 Ruth L. Kirschstein National Research Service Award from the NEI (EY029141-01) and by the University of California, San Diego, Medical Scientist Training Program T32 (GM007198-40).

